# AlphaVaR: an R framework for the statistical interpretation of AlphaGenome variant-effect predictions

**DOI:** 10.64898/2026.08.14.744948

**Authors:** Karim Marhaba, Carlo Maj, Johannes Schumacher, Pouria Dasmeh

## Abstract

**Summary:** AlphaGenome (Google DeepMind) scores a DNA variant across thousands of functional tracks at single-base resolution, reporting both the magnitude of each predicted effect and its rarity against a genome-wide background. That volume is itself the obstacle to biological interpretation. Here we present **AlphaVaR**, an R package that gives AlphaGenome’s output a typed structure together with the statistical methods and visualizations needed to interpret it. The output schema is identical for every variant, so the same tests apply throughout it. AlphaVaR provides localization tests with multiple-testing correction and effect sizes, a specificity index measuring how far an effect concentrates on a few elements of a chosen variable, and a transparent prioritization that ranks candidates across interpretable criteria and maps each to a target gene. Results feed a plot library, reproducible reports and a code-free Shiny application. Applied to rs1427407, the lead common variant for fetal-haemoglobin level, AlphaVaR recovers the established biology of the *BCL11A* erythroid enhancer.

**Availability and implementation:** https://github.com/KarimMarhaba/AlphaVaR, released under the MIT licence, R ≥ 4.2, with documentation at https://karimmarhaba.github.io/AlphaVaR/. The released version is archived at Zenodo (doi:10.5281/zenodo.21939265); the AlphaGenome scores analysed here are archived as a separate dataset (doi:10.5281/zenodo.21920988), and the scripts that regenerate every figure and reported number are in the repository (Supplementary Section S5).

**Supplementary information:** Supplementary data are available online.

## 1 Introduction

AlphaGenome (Avsec *et al*. 2026) reads up to one megabase of DNA and predicts functional genomic tracks across eleven modalities, among them chromatin accessibility, histone marks, transcription-factor binding, transcription and splicing, at single-base resolution. For every track it returns two numbers: a raw score for the physical magnitude of the predicted change, and a quantile score giving the percentile rank of that magnitude within a genome-wide background of common variants.

The same richness is what makes interpretation difficult. A single queried variant yields thousands of scored tracks, each annotated by modality, tissue, assay and finer descriptors, and returned in a long variant scoring CSV with no integrated way to combine these tracks into a variant-level readout (Supplementary Fig. S1). Identifying which tracks carry signal above the background is a prerequisite for a reproducible, interpretable analysis of AlphaGenome output.

The official AlphaGenome software (Google DeepMind 2025) provides a Python interface for running the model and a library for plotting individual tracks, and community projects add screening and language-model utilities.

Here we present AlphaVaR, an R package for the biological interpretation of AlphaGenome output. It imports the variant scoring CSV into a typed object and applies statistical tools across the whole output, turning its predictions into hypotheses that can be tested and reported. We demonstrate the application of AlphaVaR to the case of rs1427407, the lead common variant for fetal-haemoglobin level. AlphaGenome scores one alternate allele per run, so AlphaVaR groups the variant’s two alternate alleles into a single allelic series, and we follow how it reads the variant’s modality, tissue, regulators and target gene.

## 2 Implementation

### A typed object for the output

AlphaVaR imports an AlphaGenome output into an S3 object, the AlphaVarSet, which holds the full long-format variant scoring CSV together with a record of every result computed on it and of how the data was filtered (Figure 1A). The import function (`av_import_csv()`) keeps all original columns and compresses repeated metadata to factors, so even an output with tens of thousands of tracks stays compact in memory. A single AlphaVarSet can hold more than one variant at once, including the distinct alternate alleles at one position, each keyed by its `variant_id,` so an allelic series such as the T>C and T>G alleles of rs1427407 is imported into one object and analysed together. The active score, physical effect or rarity, is switched with the score-setting function (`av_set_active_score()`), and any cached result is discarded whenever the data or the active score changes, so a statistic can never be read against the wrong data. The import costs 0.33 s for the 71,420 tracks of the case study, and no analysis run on the object afterwards takes longer than two seconds, on a cohort of 3.6 million tracks either (Supplementary Section S3).

**Figure 1.**
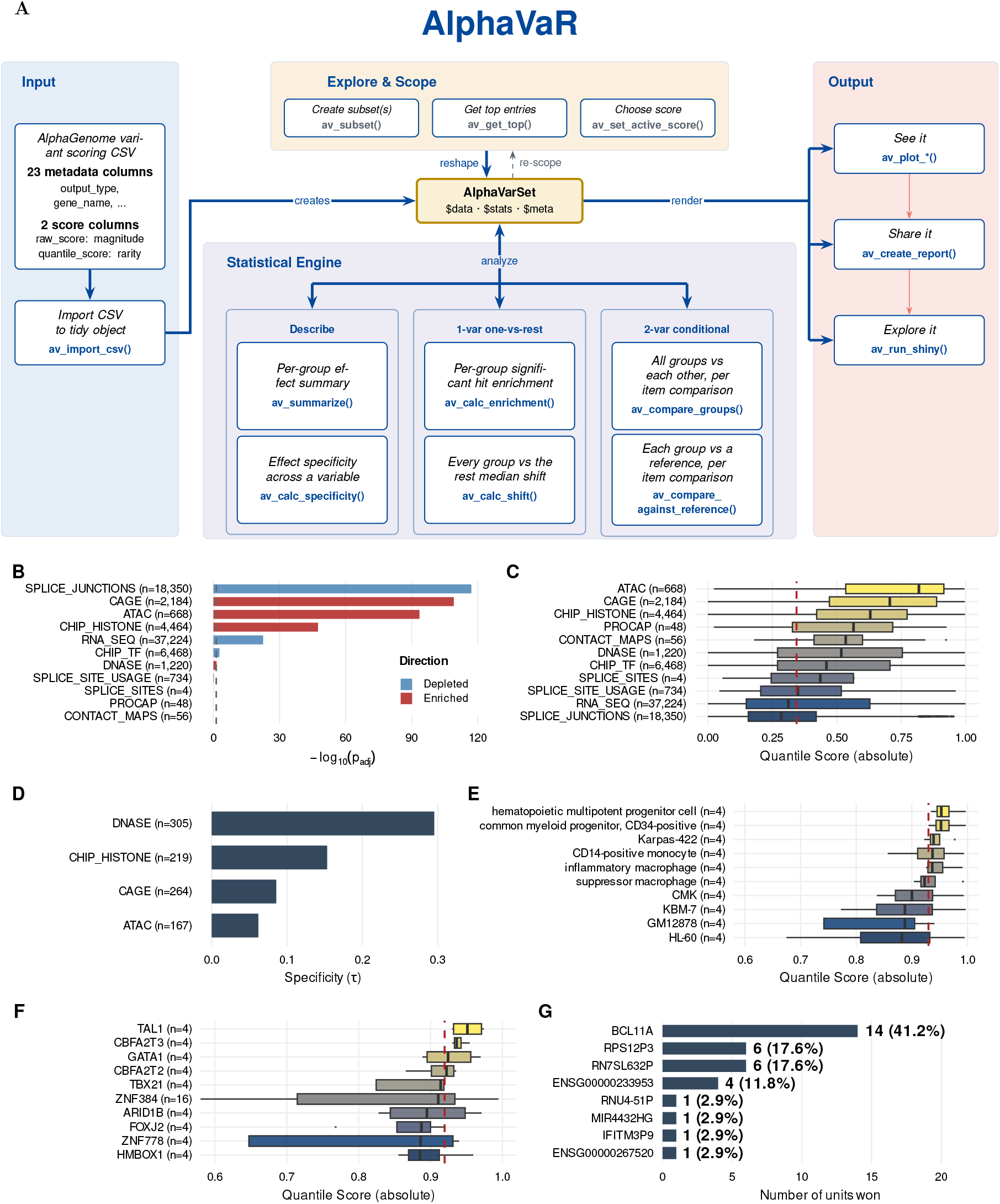
AlphaVaR and its application to rs1427407 (*BCL11A* erythroid DHS +62 enhancer, alleles T>C and T>G, hg38). **(A)** Workflow: an AlphaGenome output is imported into a typed AlphaVarSet; one set of statistical tools operates on any column of the variant scoring CSV; results feed a plot library, reproducible reports and a code-free Shiny application. **(B)** Modality enrichment (Fisher’s exact test, top-5% rarity): ATAC leads (OR = 18), splice junctions are depleted (OR = 0.009). **(C)** Absolute quantile score per modality. **(D)** Specificity (*τ*) across biosamples, among the enriched modalities: DNase is the most tissue-specific (*τ* = 0.30). **(E)** The ten top-scoring DNase biosamples, all blood-forming. **(F)** Transcription-factor ranking (descriptive, all tissues) over 751 profiled factors. **(G)** Top gene per hematopoietic tissue: *BCL11A* leads in 14 of 34. In panels B, C, E and F, *n* is the number of tracks per group; in panel D it is the number of biosamples over which *τ* is computed; in panel G the value beside each bar is the number of tissues in which the gene ranks first.

### Finding significant, non-random effects

A variant scoring CSV file contains thousands of tracks. To separate signal from noise, AlphaVaR tests each value of a chosen column against the rest. A one-vs-rest Fisher’s exact test (`av_calc_enrichment()`) asks whether extreme scores are over-represented in a group; rank-based shift tests (`av_calc_shift(), av_compare_groups()`) ask whether a group’s score distribution differs from the background, using Wilcoxon–Mann–Whitney for two groups and Kruskal–Wallis for more. All p-values are Benjamini–Hochberg corrected (Benjamini and Hochberg 1995), and every result reports an effect size with its p-value (odds ratio, signed rank-biserial *r*, or *ε*^2^): pooling thousands of tracks lets trivially small effects reach significance, so the effect size gauges both the magnitude and the direction (up- or down-regulation) that the p-value alone does not convey. The same tests run unchanged on any column, so the choice of column sets the question.

### Measuring how specific an effect is

A complementary question is how specific a variant effect is across the elements of a chosen variable. To this end, AlphaVaR computes the specificity index *τ* of Yanai *et al*. (2005) with `av_calc_specificity()`. It is bounded on [0, 1] and comparable across columns of different size, where *τ→* 0 is a uniform effect and *τ →* 1 an effect concentrated on a single element. A variant can look broad on a *τ* pooled over all modalities yet be tissue-specific within the one modality that reports the element’s activity (Figure 1D).

### Prioritizing candidate variants

When several candidate variants share a locus, the practical question is which variant is most likely to drive the regulatory effect. AlphaVaR separates this task into a target-gene assignment and a transparent ranking, and does not collapse the criteria into one synthetic score. `av_map_targets()` assigns each variant a target gene in two tiers. Tier 1 (Direct) names the highest-scoring gene among the gene-annotated tracks that pass the rarity threshold, so the call follows the predicted functional signal, and the full qualifying set is retained in `all_target_genes`. Tier 2 (Distal) is a labelled fallback for variants with no gene-annotated track above the threshold: it reports the nearest gene by the distance to the transcription start site and marks the variant as lacking direct evidence. Distance is recorded as information and never weights the ranking (Equation (1)). `av_prioritize()` then ranks variants on *K* independently interpretable criteria (three by default), with no learned or hand-set weighting: the strongest absolute per-track effect (magnitude), the tissue specificity *τ*, and the number of tracks reaching the rarity threshold (evidence breadth). Each criterion is mapped to its within-cohort percentile rank *r*_*k*_ *∈* [0, 1] and the summary rank is their mean,

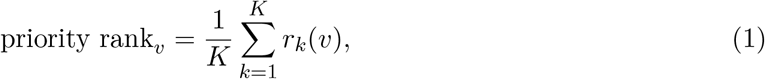

where *K* is the number of criteria and *r*_*k*_(*v*) the within-cohort percentile rank of variant *v* on criterion *k*. The mean rank is reported alongside the Pareto front, the set of variants that no other variant equals or beats on every criterion while beating it on at least one (Supplementary Section S2). A variant that leads on one criterion but falls mid-field on the others has a moderate mean rank and still lies on the front, so the two quantities are read together.

### Visualization, interactivity and reporting

Nine plotting functions render distribution, enrichment, specificity, locus and prioritization views under a shared theme. To make the pipeline usable without writing code, `av_run_shiny()` launches an interactive dashboard that exposes the full pipeline, including a causal-prioritization view that separates the slow aggregation and gene-annotation step from a fast ranking step that recomputes without rerunning the pipeline. To keep an analysis reproducible, `av_create_report()` assembles a report from user-selected analysis blocks.

### Alt text

Seven-panel figure. **(A)** Schematic of the AlphaVaR workflow. An AlphaGenome variant scoring CSV, labelled with its 23 metadata and 2 score columns, is imported into a central AlphaVarSet object; a box above it holds the subsetting and score-selection functions, a box below it groups the statistical engine into descriptive, one-vs-rest and two-variable conditional tests, and arrows lead from the object to a plotting, a reporting and a Shiny output. **(B)** Horizontal bar chart of modality enrichment, x-axis*−* log_10_ adjusted p-value, bars coloured by direction: splice junctions give the longest depleted bar, CAGE, ATAC and histone marks the longest enriched ones. **(C)**Horizontal boxplots of absolute quantile score per modality on a 0-to-1 axis, ordered from ATAC at the top to splice junctions at the bottom, with a dashed line at the global median. **(D)** Horizontal bar chart of the specificity index *τ* across biosamples for the four enriched modalities; DNase gives the longest bar, ATAC the shortest. **(E)** Horizontal boxplots of the ten highest-scoring DNase biosamples on a zoomed 0.6-to-1 axis, all of them blood-forming, led by a hematopoietic multipotent progenitor cell. **(F)** The same boxplot form for the ten highest-ranking transcription factors, led by TAL1, CBFA2T3, GATA1 and CBFA2T2. **(G)** Horizontal bar chart counting the hematopoietic tissues in which each gene ranks first, each bar labelled with its count; *BCL11A* gives by far the longest bar, at 14 tissues.

## 3 Application: rs1427407 in the *BCL11A* erythroid enhancer

We applied AlphaVaR to rs1427407 (chr2:60,490,908, hg38; alleles T>C and T>G), the lead common variant for fetal-haemoglobin (HbF) level (Bauer *et al*. 2013). *BCL11A* encodes a zinc-finger transcriptional repressor that silences the *γ*-globin genes and so represses HbF in adult erythroid cells. The variant marks the +62 DNase-hypersensitive site of the gene’s erythroid enhancer, whose disruption reactivates HbF and is the mechanism of the approved CRISPR therapy for sickle-cell disease and *β*-thalassaemia (Canver et al. 2015, Frangoul et al. 2021). Because this biology is known, the variant is a positive control. The two alleles were scored across 11 modalities and up to 714 biosamples, giving 71,420 track predictions (Figure 1). In the following sections, we first introduce the biological questions that AlphaGenome predictions can address, then answer each of them with AlphaVaR.

### Which modalities carry the effect?

We first tested, across modalities, where extreme scores were over- or under-represented. At the top-5% rarity default, chromatin and active-regulatory modalities were strongly enriched (Figure 1B; ATAC odds ratio 18, log_2_ OR 4.2; CAGE and histone marks following at log_2_ OR 3.2 and 2.0), while splice junctions were depleted (OR 0.009), and these enriched channels also carried the highest absolute quantile scores per track (Figure 1C). Summarized per modality, the signed raw score was a gain in all eleven modalities, with the same direction in 96% of ATAC tracks, consistent with the variant increasing the regulatory activity.

### In which tissue/cell type does it act?

Specificity depends on the channel it is read in: among the enriched modalities *τ* ranges from 0.06 (ATAC) to 0.30 (DNase; Supplementary Table S3). DNase, the assay that defines this hypersensitive site, carried the most tissue-specific signal (Figure 1D). Reading tissues within DNase, the ten highest-scoring biosamples were all blood-forming, led by a hematopoietic multipotent progenitor (Figure 1E); on the raw score the channel sharpened further (*τ* 0.84, top tissue a CD34-positive common myeloid progenitor). The effect was erythroid and hematopoietic, recovered once the specificity was read in the right modality.

### Which regulators drive it?

Having placed the variant’s effect in accessible chromatin within the hematopoietic lineage, we then ranked 751 profiled transcription factors by their effect on TF-binding tracks (Figure 1F). The erythroid master-regulator complex topped the ranking, TAL1, CBFA2T3 (ETO2), GATA1 and CBFA2T2, and rs1427407 lies in the GATA1/TAL1 occupancy peak of the enhancer (Bauer et al. 2013, Canver et al. 2015). GATA1 and TAL1 are profiled on only four tracks each, in K562 cells, so a minimum-coverage filter would drop them and place a broadly profiled factor first. The ranking was therefore reported descriptively, without such a filter.

### Which gene is the target?

The 1 Mb scored interval contains 23 annotated genes, and only three of them are protein-coding, namely *BCL11A, PAPOLG* and *REL*. `av_map_targets()` returned *BCL11A* as the Tier-1 target for the T>C allele, robust across mean, median and max aggregation; for T>G only max aggregation returned *BCL11A*, so the aggregation choice was reported alongside the call. Read per hematopoietic tissue, *BCL11A* was also the most frequent top gene (14 of 34 tissues; Figure 1G), whereas pooling all tissues placed it sixth, behind neighbouring pseudogenes.

Across the four steps, AlphaVaR reconstructed the established biology of rs1427407 from Al-phaGenome’s predictions: a non-coding enhancer with increased predicted activity, erythroid in lineage, driven by the GATA1/TAL1 complex, targeting *BCL11A*. The analysis also returned negative results where the biology predicts them, with splice junctions depleted and specificity low in most channels.

## 4 Discussion

When handed an AlphaGenome output, the question is rarely whether a variant has some predicted effect, as predictions are distributed across numerous modalities, assays and biosamples. Rather, the challenge is to identify which signals are reproducible across related contexts and statistically compelling enough to warrant biological interpretation and experimental follow-up. The ecosystem so far leaves this step to the user. AlphaVaR fills it with a statistical layer carrying multiple-testing control and effect sizes, and with an interface that opens the whole pipeline without the need for programming. It complements single-score variant-effect predictors such as CADD, REVEL and AlphaMissense, which reduce each variant to a scalar and do not model the modality-by-tissue-by-track structure that AlphaGenome produces. Existing tools run AlphaGenome and visualize its tracks (Abir 2025) but do not test that output statistically, so there is no established baseline for functional prioritization to benchmark against.

AlphaVaR performs variant-level epigenomic interpretation and conditional prioritization within a locus. It does not infer phenotype-level causality. Which tissue mediates a trait, or which mechanism drives it, needs external evidence such as GWAS–tissue colocalization or eQTL fine-mapping. Nevertheless, AlphaVaR prioritizes the predicted molecular consequences of GWAS variants across tissues, cell types and modalities, functional evidence that complements statistical fine-mapping and variant prioritization.

Several limitations follow from our approach. A single variant scored over thousands of tracks invites pseudo-replication: cross-track p-values track assay coverage more than effect, so the effect sizes carry the interpretation. Specificity measures magnitude concentration and is diluted by pooling across modalities, so the lineage is recovered only by conditioning on the signal-defining modality. The minimum-coverage guard removes small-sample artefacts but can drop genuine low-coverage regulators, so descriptive ranking suits discovery and the tests suit well-covered groups. A sequence-only model predicts binding-site disruption but not cell-type-specific function, so the predicted footprint is broader than the verified erythroid restriction, and a co-signal in a non-erythroid line is expected. Many tracks saturate near the quantile ceiling, so the raw score separates tissues and factors better. Assigning a Tier-2 target gene by transcription-start-site distance is a deliberate simplification: regulatory targeting need not track linear distance, and robust SNP-to-gene mapping remains an open problem. The priority rank orders candidates within a locus and carries no significance interpretation. Finally, we report no quantitative benchmark against single-score predictors such as CADD, REVEL or AlphaMissense, because these reduce a variant to one scalar and leave the modality, tissue and track structure that AlphaVaR ranks on unresolved.

Future work will track AlphaGenome model updates, develop a supervised approach to variant prioritization against experimentally validated targets, and pursue submission to Bioconductor.

