## Supplementary Material for "AlphaVaR: an R framework for the statistical interpretation of AlphaGenome variant-effect predictions"

### Contents


- **S1** The AlphaGenome output that AlphaVaR reads (Figure S1)
- **S2** The Pareto front used by the prioritization (Equations S1, S2)
- **S3** Runtime and memory (Tables S1, S2)
- **S4** Per-modality tissue specificity for rs1427407 (Table S3)
- **S5** Reproducing the figures and results (Tables S4, S5)
- References


The table underlying Section S4 is printed here in full, so there is no separate supplementary data file.

### S1 The AlphaGenome output that AlphaVaR reads

Figure S1 shows the raw material the package works on: an excerpt of the variant scoring CSV for rs1427407, and the number of tracks each modality contributes to it. It is the concrete form of the interpretation problem the manuscript opens with. One queried variant produces 71,420 scored rows in 25 columns, the file carries no ranking of its own, and the track budget differs by four orders of magnitude between modalities, so a signal in a small channel is easily outweighed by the volume of a large one.

**A One AlphaGenome output: 71,420 rows x 25 columns, no built-in ranking**

| variant_id | scored_interval | output_type | Assay title | biosample_name | biosample_type | data_source | gene_name | +15 more | raw_score | quantile_score |
|---|---|---|---|---|---|---|---|---|---|---|
| chr2:60490908:T>C | chr2:59966621-61015197:. | ATAC | ATAC-seq | T-cell | primary_cell | encode | | … | 0.0845 | 0.9524 |
| chr2:60490908:T>C | chr2:59966621-61015197:. | DNASE | DNase-seq | neuronal stem cell | in_vitro_differen… | encode | | … | 0.0068 | 0.2267 |
| chr2:60490908:T>C | chr2:59966621-61015197:. | CHIP_TF | TF ChIP-seq | osteoblast | primary_cell | encode | | … | -0.0126 | -0.6618 |
| chr2:60490908:T>C | chr2:59966621-61015197:. | CHIP_HISTONE | Histone ChIP-seq | neuronal stem cell | in_vitro_differen… | encode | | … | -0.0131 | -0.6618 |
| chr2:60490908:T>C | chr2:59966621-61015197:. | CAGE | LQhCAGE | mesothelial cell | primary_cell | fantom | | … | 0.0033 | 0.5287 |
| chr2:60490908:T>C | chr2:59966621-61015197:. | RNA_SEQ | polyA plus RNA-s… | neuronal stem cell | in_vitro_differen… | encode | PAPOLG | … | -0.0002 | -0.3832 |
| chr2:60490908:T>C | chr2:59966621-61015197:. | SPLICE_JUNCTIONS | polyA plus RNA-s… | neuronal stem cell | in_vitro_differen… | encode | BCL11A | … | 0.0177 | 0.6916 |
| chr2:60490908:T>C | chr2:59966621-61015197:. | SPLICE_SITE_USAGE | polyA plus RNA-s… | neuronal stem cell | in_vitro_differen… | encode | BCL11A | … | 0.0039 | 0.5744 |
| chr2:60490908:T>C | chr2:59966621-61015197:. | CONTACT_MAPS | Micro-C | H1-hESC | cell_line | 4dnucleome | | … | 0.0009 | 0.5801 |
| chr2:60490908:T>C | chr2:59966621-61015197:. | PROCAP | PRO-cap | A673 | cell_line | encode | | … | 0.0096 | 0.5844 |
| ⋮ | ⋮ | ⋮ | ⋮ | ⋮ | ⋮ | ⋮ | ⋮ | | ⋮ | ⋮ |

**B Tracks per modality**

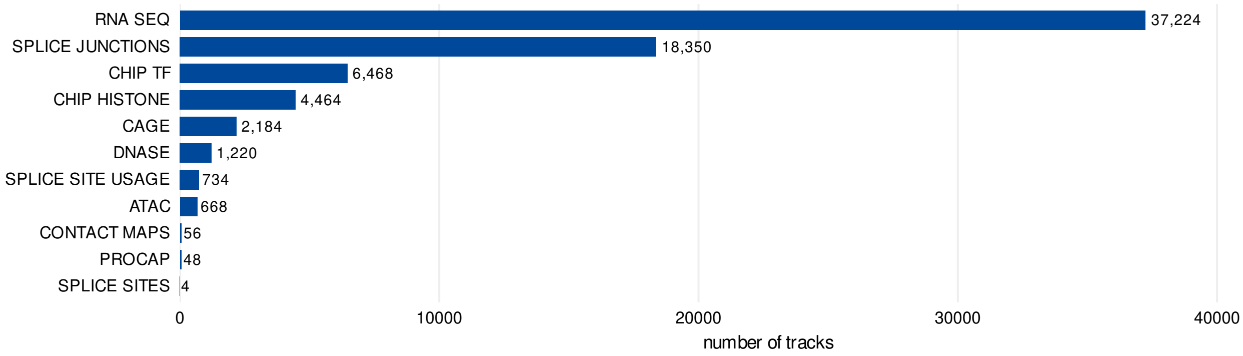


Figure S1: The raw AlphaGenome output for a single variant, before AlphaVaR. **(A)** Ten rows of the variant scoring CSV for rs1427407, one per modality, showing 10 of its 25 columns. The two score columns are pinned to the right; the remaining 15 metadata columns are collapsed into the ellipsis column. **(B)** Track counts per modality range from 4 (splice sites) to 37,224 (RNA-seq), so a signal in a small channel is easily outweighed by the volume of the large ones.

### S2 The Pareto front used by the prioritization

`av_prioritize()` reports two quantities per variant: the mean percentile rank of Equation (1) of the manuscript, and whether the variant lies on the Pareto front. The front is defined as follows. For a cohort $V$ of candidate variants scored on $K$ criteria, with $r_k(v)$ the within-cohort percentile rank of variant $v$ on criterion $k$, a variant $u$ dominates a variant $v$ when it is at least as good on every criterion and strictly better on at least one,

$$u \succ v \iff r_k(u) \geq r_k(v)\ \forall k \in \{1, \ldots, K\}\ \wedge\ \exists\, k : r_k(u) > r_k(v), \tag{S1}$$

and the front is the set of variants that no other variant dominates,

$$\mathcal{P} = \{\, v \in V : \neg\, \exists\, u \in V \text{ with } u \succ v \,\}. \tag{S2}$$

The two quantities answer different questions and are read together. The mean rank is a single ordering, in which a variant that is merely consistent across criteria can outrank a variant that leads on one; the front keeps every variant that no other variant beats outright, so a specialist candidate stays visible. Neither quantity uses a learned or hand-set weighting, and neither carries a significance interpretation.

### S3 Runtime and memory

Table S1 gives the wall-clock time and the additional memory of each analysis step on the full rs1427407 table, and Table S2 how both grow with cohort size. All figures are medians of 3 runs on Intel Core Ultra 5 125U (14 cores, 31 GB RAM), R 4.6.1, single-threaded.

No step of a single-variant analysis takes longer than 0.48 s. Reading the 20 MB CSV costs 0.14 s, and the statistical steps that follow cost between 0.03 and 0.48 s each. Because the import is paid once and not per question asked of the object, an analysis session stays interactive, which is the condition the Shiny application depends on.

Table S2 scales the same pipeline to cohorts. Memory grows in proportion to the number of scored tracks, wall-clock more slowly: fifty times the data costs 24 times the time and 15 times the memory, because the fixed cost of grouping is amortized over more rows. The largest cohort tested, 50 variants, 3,571,000 tracks, passes through enrichment, specificity and summarization in 1.67 s using 1.3 GB above the imported table. A locus of a few hundred variants is therefore a laptop-scale analysis; the runtime of a study is set by the AlphaGenome query that produces the scores, not by their interpretation.

Table S1: Wall-clock time and additional memory per analysis step on the full rs1427407 table (71,420 scored tracks). Memory is the peak allocation the step adds to the session, not the total footprint.

| Step | What it computes | Seconds | Memory (MB) |
|---|---|---|---|
| `av_import_csv` | 71,420 rows x 25 columns from CSV | 0.14 | 33 |
| `av_calc_enrichment` | Fisher one-vs-rest over 11 modalities | 0.03 | 28 |
| `av_calc_specificity` | Tau per modality across biosamples | 0.03 | 38 |
| `av_summarize` | variant-level recipe, balanced by modality | 0.03 | 34 |
| `av_calc_shift` | one-vs-rest sweep over 11 modalities | 0.48 | 112 |

Table S2: Enrichment, specificity and summarization on synthetic cohorts built by stacking the rs1427407 table under distinct variant identifiers, which preserves the modality and biosample composition of a genuine AlphaGenome export.

| Cohort | Seconds | Memory (MB) |
|---|---|---|
| 1 variant, 71,420 tracks | 0.07 | 86 |
| 5 variants, 357,100 tracks | 0.19 | 239 |
| 20 variants, 1,428,400 tracks | 0.61 | 666 |
| 50 variants, 3,571,000 tracks | 1.67 | 1297 |

### S4 Per-modality tissue specificity for rs1427407

Table S3 reports the specificity index $\tau$ computed per modality across biosamples, on both score domains, for the lead variant rs1427407. The index follows Yanai *et al.* (2005). It is the per-modality scan behind panel D of Figure 1. In the table, $n_{\text{active}}$ counts the biosamples whose score passes the rarity threshold, $n_{\text{biosamples}}$ is the number of distinct biosamples over which $\tau$ is computed, and the top biosample is the one carrying the largest share of the signal. Ten of the eleven AlphaGenome modalities appear: `SPLICE_SITES` is scored without a biosample annotation, so a specificity across biosamples is not defined for it.

$\tau$ is bounded on $[0, 1]$: $\tau \to 0$ is a uniform effect across biosamples, $\tau \to 1$ an effect concentrated on a single one. It is a descriptive metric and carries no p-value.

The two score domains answer different questions and are not interchangeable. The quantile domain measures statistical rarity against a genome-wide background, so it is the domain in which tissue specificity is interpretable; the raw domain measures the physical magnitude of the predicted change, in which the largest values follow track abundance rather than biology.

Panel D of Figure 1 restricts the comparison to the four modalities in which extreme scores are significantly over-represented for this variant (`ATAC`, `CAGE`, `CHIP_HISTONE`, `DNASE`; Fisher's exact test at the top-5% rarity default, $\log_2 \text{OR} > 0$ and $p_{\text{adj}} < 0.05$ after the correction of Benjamini and Hochberg (1995)). Within that set DNase is the most tissue-specific in the quantile domain ($\tau = 0.30$), which is the observation the case study builds on. Read over all ten modalities in Table S3 the ranking is a different one: `SPLICE_SITE_USAGE` ($\tau = 0.40$) and `CONTACT_MAPS` ($\tau = 0.31$) sit above DNase. Neither carries a significant enrichment of extreme scores for this variant (`SPLICE_SITE_USAGE` $\log_2 \text{OR} = -0.51$, $p_{\text{adj}} = 0.58$; `CONTACT_MAPS` $\log_2 \text{OR} = -0.63$, $p_{\text{adj}} = 1.0$), so a high $\tau$ there concentrates a signal that is not distinguishable from the background in the first place. That is the reason the case study conditions on the signal-defining modality before reading $\tau$, rather than taking the maximum over all channels.

Table S3: Per-modality tissue specificity for rs1427407 on both score domains.

| Modality | Domain | $\tau$ | Top biosample | $n_{\text{active}}$ | $n_{\text{biosamples}}$ |
|---|---|---|---|---|---|
| ATAC | quantile | 0.0620 | naive B cell | 118 | 167 |
| ATAC | raw | 0.3373 | HG03469 | 167 | 167 |
| CAGE | quantile | 0.0858 | bone marrow cell | 73 | 264 |
| CAGE | raw | 0.8625 | temporal lobe | 210 | 264 |
| CHIP_HISTONE | quantile | 0.1533 | CD14-positive monocyte | 52 | 219 |
| CHIP_HISTONE | raw | 0.8437 | DOHH2 | 219 | 219 |
| CHIP_TF | quantile | 0.2580 | HL-60 | 11 | 163 |
| CHIP_TF | raw | 0.5255 | K562 | 163 | 163 |
| CONTACT_MAPS | quantile | 0.3114 | GM12878 | 0 | 12 |
| CONTACT_MAPS | raw | 0.2817 | GM12878 | 0 | 12 |
| DNASE | quantile | 0.2955 | hematopoietic multipotent progenitor cell | 23 | 305 |
| DNASE | raw | 0.8372 | common myeloid progenitor, CD34-positive | 305 | 305 |
| PROCAP | quantile | 0.1880 | K562 | 0 | 6 |
| PROCAP | raw | 0.5304 | K562 | 2 | 6 |
| RNA_SEQ | quantile | 0.0559 | heart | 151 | 285 |
| RNA_SEQ | raw | 0.7594 | GM12878 | 15 | 285 |
| SPLICE_JUNCTIONS | quantile | 0.2665 | K562 | 2 | 282 |
| SPLICE_JUNCTIONS | raw | 0.5553 | K562 | 0 | 282 |
| SPLICE_SITE_USAGE | quantile | 0.4007 | thoracic aorta | 7 | 282 |
| SPLICE_SITE_USAGE | raw | 0.2856 | A375 | 0 | 282 |

### S5 Reproducing the figures and results

Every figure and every number in the manuscript is produced by a script in the `paper/` directory of the software repository (Marhaba 2026). Table S4 maps each manuscript item to its script, the AlphaVaR functions that carry the analysis, and the file it writes. The numbers the manuscript states in running text rather than in a figure are covered by `scripts/make_reported_numbers.R`, which recomputes each of them and stops with an error if any has drifted from the value the text prints. Figure 1 is assembled from two inputs: the workflow schematic in panel A and the case-study block in panels B–G are built separately and stitched into one file by `figures/figure1_workflow_case.tex`.

Table S4: Manuscript item, the script that builds it, the AlphaVaR functions it calls, and the file it writes. Script and output paths are relative to `paper/` in the repository.

| Item | Script | Calls | Output |
|---|---|---|---|
| Figure 1 (whole) | `figures/figure1_workflow_case.tex` | none (stitches the two panels below) | `figures/figure1_workflow_case.pdf` |
| Figure 1, panel A | `figures/figure1_overview.tex` | none (hand-drawn TikZ) | `figures/figure1_overview.pdf` |
| Figure 1, panels B–G | `scripts/make_figure2.R+ figure1_calls.R` | `av_import_csv`, `av_calc_enrichment`, `av_subset`, `av_calc_specificity`, `av_summarize`, `av_plot_enrichment`, `av_plot_distribution`, `av_plot_specificity` | `figures/figure2_bcl11a_case_study.pdf` |
| Figure S1 | `scripts/make_figure1_intro.R` | `av_theme_alpha` | `figures/figure1_intro.pdf` |
| Tables S1–S2 | `scripts/make_s_runtime.R` | `av_import_csv`, `av_calc_enrichment`, `av_calc_specificity`, `av_summarize`, `av_calc_shift` | `supplementary/runtime_*.csv` |
| Table S3 | `scripts/make_tau_table.R` | `av_import_csv`, `av_set_active_score`, `av_calc_specificity` | `supplementary/per_modality_tau.csv` |
| Numbers stated in the text | `scripts/make_reported_numbers.R` | `av_calc_enrichment`, `av_summarize`, `av_set_active_score`, `av_map_targets`, `av_calc_specificity`, `av_subset` | `supplementary/reported_numbers.csv` |

#### Running the scripts

The R scripts are run from the repository root, which is where `paper/scripts/_data.R` resolves the input table from:

```
Rscript paper/scripts/make_figure1_intro.R
Rscript paper/scripts/make_figure2.R
Rscript paper/scripts/make_s_runtime.R
Rscript paper/scripts/make_tau_table.R
Rscript paper/scripts/make_reported_numbers.R
```

They need the package source tree rather than the installed package, because they load it with `devtools::load_all()`, and the first of them downloads the 20 MB score table unless a local copy is already in place.

Panel A of Figure 1 is a hand-drawn schematic, and the two halves of Figure 1 are stitched together, so both steps are compiled with LaTeX instead:

```
pdflatex -output-directory paper/figures paper/figures/figure1_overview.tex
pdflatex -output-directory paper/figures paper/figures/figure1_workflow_case.tex
```

#### Code for panels B to G of Figure 1

Table S4 names the functions each script calls; the arguments decide what the functions answer. The analysis of the case study is therefore kept in its own file, `scripts/figure1_calls.R`, which `scripts/make_figure2.R` sources before it draws the panels. It is printed below in full, so the calls shown here are the calls that produced the figure. Arguments left at their default are not written out; those below are the choices the case study makes.

```r
obj <- av_import_csv(csv, verbose = FALSE)

## B: are extreme scores over-represented in one modality?
obj <- av_calc_enrichment(obj, group_by = "output_type")

## C: the magnitude per modality is drawn from `obj` itself, so no call of
## its own appears here.

## D: tissue specificity, read only in the modalities the Fisher test marks
## as enriched. CHIP_TF is excluded although it is a chromatin channel,
## because its enrichment is negative and significant (log2OR -0.62,
## p_adj 0.0019); SPLICE_SITES because its positive log2OR rests on four
## tracks.
SIG <- c("ATAC", "CAGE", "CHIP_HISTONE", "DNASE")
sig <- av_calc_specificity(av_subset(obj, output_type %in% SIG),
                           per = "output_type",
                           across = "biosample_name", metric = "both")
```

```r
## E: which biosamples carry the DNase signal, and the ten leading ones
dnq <- av_calc_specificity(av_subset(obj, output_type == "DNASE"),
                           per = "variant_id",
                           across = "biosample_name", metric = "both")
dnq <- av_summarize(dnq, group_by = "biosample_name")
dq  <- dnq$stats$summary[["biosample_name"]]
dq  <- dq[order(-dq$median_score), ]

## F: which transcription factors, and the ten leading ones
tf  <- av_summarize(av_subset(obj, output_type == "CHIP_TF"),
                    group_by = "transcription_factor")
tfb <- tf$stats$summary[["transcription_factor"]]
tfb <- tfb[order(-tfb$median_score), ]

## G: the target gene, read across the hematopoietic biosamples only
hema_rx <- paste0(
  "blood|lymph|B cell|T cell|monocyt|macrophag|erythro|myelo|leukem|",
  "lymphoblast|dendritic|mast|eosinophil|basophil|neutrophil|megakaryo|",
  "hematopoiet|mononuclear|reticulocyte|GM1|HG0|K562|Karpas|THP|marrow|",
  "killer|thymocyte|HL-60|OCI|Jurkat|Langerhans|progenitor")
HEMA <- unique(obj$data$biosample_name[
  grepl(hema_rx, obj$data$biosample_name, ignore.case = TRUE)])
gen <- av_subset(obj, output_type == "RNA_SEQ" & biosample_name %in% HEMA)
gen <- av_calc_specificity(gen, per = "biosample_name",
                           across = "gene_name", metric = "both")
```

#### Input data

The scripts read one file: the AlphaGenome (Avsec *et al.* 2026) variant-effect scores for rs1427407 (GRCh38, 1 Mb context, alleles T>C and T>G; 71,420 scored tracks across 11 modalities). At 20 MB it is deliberately kept out of the git repository and out of the package tarball, and is archived at Zenodo instead (Marhaba *et al.* 2026).

`paper/scripts/_data.R` resolves it in a fixed order (local working copy, then user cache, then download from the archived record) and verifies the download against a recorded SHA-256 checksum, so a truncated transfer or a silently replaced record fails loudly rather than producing different figures. The scripts call `bcl11a_scores()` from that file rather than hardcoding a path, so the figures reproduce from the archived deposit alone.

The table itself was produced by `paper/scripts/generate_bcl11a_scores.py`, which queries the AlphaGenome API for the two alternate alleles and writes the CSV the package reads.

The same table ships with the package, gzipped to 1.9 MB, as `inst/extdata/bcl11a_rs1427407_scores.csv.gz`, so the vignette reproduces the reported values without network access. A seeded stratified subset of it, preserving all 11 modalities and the full 25-column schema, ships alongside and backs the package examples and the test suite, where a small file keeps runtimes short; because it caps each modality at a fixed number of rows, it is unsuitable for the enrichment tests, whose results depend on the relative sizes of the modalities.

#### Software environment

Table S5: Versions used to render this supplement.

| Component | Version |
|---|---|
| OS | Ubuntu 24.04.4 LTS |
| R | 4.6.1 |
| AlphaVaR | 0.0.0.9000 |
| Repository commit | 1c070ff (working tree modified) |
| dplyr | 1.2.1 |
| tidyr | 1.3.2 |
| ggplot2 | 4.0.3 |
| patchwork | 1.3.2 |
| GenomicRanges | 1.64.0 |

The package requires R $\geq$ 4.2. Its full dependency set, with the versions the release was checked against, is recorded in `DESCRIPTION`; the continuous-integration workflow in `.github/workflows/` runs `R CMD check`, including the test suite, on a clean machine for every change. The package documentation carries a worked end-to-end vignette on the same table, at `https://karimmarhaba.github.io/AlphaVaR/`.

The version that produced everything reported here is archived at Zenodo (Marhaba 2026), and that record, not the repository head, is what the numbers in the manuscript belong to. No step of the pipeline is stochastic, so no seed is set anywhere in the scripts; the one random draw in the project is the seeded stratified subset that backs the package examples and the test suite, and it enters no figure and no reported number.
